# dbDEMC 3.0: Functional Exploration of Differentially Expressed miRNAs in Cancers of Human and Model Organisms

**DOI:** 10.1101/2022.02.10.479911

**Authors:** Feng Xu, Yifan Wang, Yunchao Ling, Chenfen Zhou, Haizhou Wang, Andrew E. Teschendorff, Yi Zhao, Haitao Zhao, Yungang He, Guoqing Zhang, Zhen Yang

## Abstract

microRNAs (miRNAs) are important regulators in gene expression. The deregulation of miRNA expression is widely reported in the transformation from physiological to pathological state of cells. A large amount of differentially expressed miRNAs (DEMs) have been identified in various human cancers by using high-throughput technologies, such as microarray and miRNA-seq. Through mining of published researches with high-throughput experiment information, the database of differentially expressed miRNAs in human cancers (dbDEMC) was constructed with the aim of providing a systematic resource for the storage and query of the DEMs. Here we report an update of the dbDEMC to version 3.0, containing two-fold more data entries than the previous version, now including also data from mouse and rat. The dbDEMC 3.0 contains 3,268 unique DEMs in 40 different cancer types. The current datasets for differential expression analysis have expanded to 9 generalized categories. Moreover, the current release integrates functional annotations of DEMs obtained from experimentally validated targets. The annotations can greatly benefit integrative analysis of DEMs. In summary, dbDEMC 3.0 provides a valuable resource for characterizing molecular functions and regulatory mechanisms of DEMs in human cancers. The dbDEMC 3.0 is freely accessible at https://www.biosino.org/dbDEMC.

## Introduction

Since the first discovery of microRNAs (miRNAs) at the beginning of this century, this class of small non-coding RNA has received extensive attention [1]. As an important gene expression regulator acting at the post-transcriptional level, studies have disclosed the critical role of miRNAs in targeting mRNAs for the degradation or translational repression [2]. Till now, a total of 2,654 miRNAs have been identified in the human genome according to the latest version of miRBase database [3]. Significant researches on miRNAs have dramatically expanded our understanding about gene regulatory network and its roles in physiological and pathological conditions, such as in broad spectrums of biological processes including cell cycle, cell proliferation, differentiation, apoptosis and cellular signaling [4, 5]. Owing to the biological significance of miRNAs, alterations of the miRNAs are linked to the development of many diseases including the cancer [6]. Differentially expressed miRNAs (DEMs) are widely reported to hold great value in the diagnosis or prognosis as well as treatment targeting for cancer research [7]. The potential use of circulating miRNAs in serum, plasma and other body fluids as non-invasive cancer biomarkers have also been thoroughly investigated [8].

Given the important functions of miRNA in cancer development, several on-line resources have been built for warehousing information of cancer-related miRNAs, such as the HMDD [9], miRCancer [10], and OncomiRDB [11]. With the development of high-throughput techniques such as microarray and miRNA-seq, large amount of cancer DEMs were identified from miRNA profiling data each year. However, these valuable data were scattered in the vast literatures and it is of great necessity to catalogue them in a favorable way, thus to provide integrative tools for the effective utilization and systematic investigation. With this aim, we developed the initial database of differentially expressed miRNAs in human cancers (dbDEMC) in 2010 [12] and further updated in 2017 [13]. To our knowledge, dbDEMC is the only working repository currently available for storing DEMs from de-novo analysis of high-throughput profiling data in human cancers, characteristic with miRNANome data in various types of cancer. It greatly facilitates the efforts to excavate cancer associated miRNAs and investigate their roles in pathological processes of cancer. While the database could have been much more useful had there been more high-quality data included.

In recent years, cancer quantitative miRNA profiling data has been increasing at an unprecedented rate, and given the success of dbDEMC 2.0, this motivates an update of this database. Here we introduce dbDEMC 3.0, a significantly expanded version of this database. This update incorporates a substantial amount of new data. Besides the human data, we have also incorporated the miRNA expression profiling data of mouse and rat. A total of 403 datasets of high-throughput miRNA expression encompassing 40 cancer-types, with the results of 807 differential expression analyses, have been included. The present update is nearly doubling the data amount over the previous version. In addition to the expanded data volume, the content of the database has also been enriched. The new database firstly recorded the experimentally validated DEMs targets and also their enrichment analysis on Gene Ontology (GO) and Kyoto Encyclopedia of Genes and Genomes (KEGG) pathways. In this way, we provided the functional annotation of DEMs for various cancers. Also, the web interface of the database was refined for a better visualization of the aforementioned data. Taken together, the dbDEMC 3.0 is a comprehensive resource to systematically characterize the function of DEMs in human cancers as well as other model organisms.

## Data collection and processing

### Data collection

To compile the datasets, we used the keywords ‘cancer’, ‘tumor’, ‘carcinoma’ and ‘neoplasm’, in combination with ‘microRNA’ or ‘miRNA’ to conduct an exhaustive search for the microarray based miRNA expression profiles in Gene Expression Omnibus (GEO) [14] and ArrayExpress [15], and further for miRNA-seq based miRNA expression profiles in Sequence Read Archive (SRA) [16]. While the miRNA profiles of mouse and rat were rapidly accumulating, we also incorporated the miRNA data of the two model organisms in the current update. In addition, we also appended the new miRNA profiling data from TCGA that was released since the last update of dbDEMC 2.0. All the involved data were published as of June 2021. The data records were manually reviewed and evaluated rigorously to guarantee that only high-quality data sets were included. To ensure analysis reliability, we required at least three biological replicates of samples each condition (for both case and control) as usual.

### Data processing

For miRNA profiling data sets based on microarray, we used the same protocol as that of dbDEMC 2.0 to identify the DEMs [13]. Briefly, the expression values were logarithmically transformed (base 2) and quantile normalized. Then the limma (Linear Models for Microarray Data) package was applied to select miRNAs whose mean expression level is significantly different between case and control samples with FDR value < 0.05.

For miRNA-seq based profiling data obtained from SRA database, we downloaded the SRA files of raw sequence reads and converted them into FASTQ format using the fastq-dump of SRA Toolkit. Here we included only the data produced on Illumina systems (Genome Analyzer I, II, IIx, HiSeq 1000, HiSeq 2000, HiSeq 2500, HiSeq 4000, NextSeq, MiSeq). The involving miRNA-seq data was analyzed by using QuickMIRSeq toolkit [17]. This toolkit utilizes the Cutadapt to remove sequence adapters and performs quality control [18]. We collected detailed information of DNA adapters of different miRNA-seq libraries from public resources to guarantee that the adapters can be properly trimmed from the raw reads (Table S1) [19]. The clean reads were then aligned to the reference genome by using Bowtie [20], and miRDeep2 was used to obtain count tables of aligned reads for miRNA quantification [21]. The reads count table was further normalized by using limma-voom [22], and DEMs were then identified. For the datasets obtained from TCGA, we directly used the prepared reads count data for further analysis [23].

Experimental validation of DEMs in low-throughput methods (such as real-time PCR and northern blot, etc.) were manually collected from the original papers. These types of information were carefully formatted and integrated into our update.

### Functional annotation

For each obtained DEMs set, we collected the experimentally validated targets by using multiMiR [24], which integrate miRNA target data from TarBase [25] and miRTarBase [26]. Then we performed the enrichment analysis of the DEMs targets on GO terms and KEGG pathways by using clusterProfiler package to facilitate the study of context-dependent miRNA functional mechanisms [27]. Enriched GO terms and KEGG pathways were selected where adjusted P value < 0.05. The data collection and curation procedure for dbDEMC 3.0 is shown in **Figure 1**.

**Figure 1.**
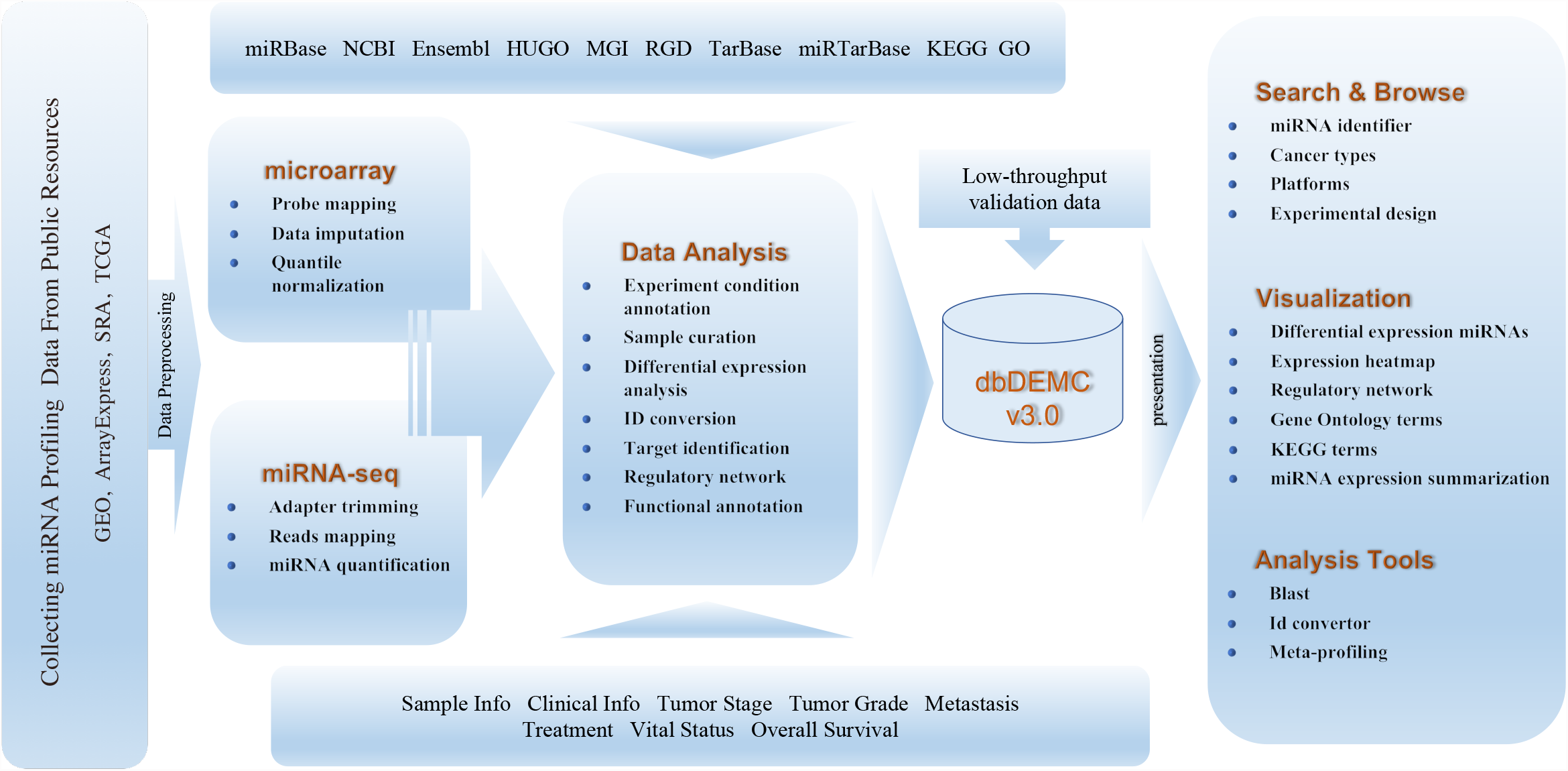
Schematic illustration of the data collection and architecture of the dbDEMC 3.0.

### Database construction

All the data in dbDEMC 3.0 was managed by using MongoDB. The dynamic web interface was developed using JSP and JavaScript. Data visualization was achieved through the tools of vue, jquery, and Echarts, Elasticsearch was used for search engine. The database was developed by Spring Boot framework. Apache Tomcat web server was used for the http server. All the information in dbDEMC 3.0 is freely available to the public domain through https://www.biosino.org./dbDEMC.

### miRNA cluster annotation

A miRNA cluster is defined as set of miRNAs which located adjacent genomic regions in the same or opposite orientation and not separated by other transcriptional units. miRNAs within a cluster are thought to be regulated by common factors and involved in same signaling pathways. According to Kabekkodu SP et al., among 1881 precursor miRNAs of human origin annotated in miRBase, 468 of which can be attributed to 153 clusters [28]. Here we obtained these data about miRNA clusters and annotated mature miRNAs by using annotation file from miRBase. Finally, a total of 688 (22.8%) mature miRNAs from 143 clusters were annotated in the human genome. Analysis of homogeneous dysregulation pattern of miRNA clusters in cancer For systematic study of co-dysregulation pattern of miRNA clusters in human cancers, we considered all miRNAs associated with specific cancer. miRNAs not belonging to any cluster and clusters of which at least half the members are not associated with any cancer were discarded. To avoid potential bias introduced by different expression platforms, here we only use the results obtained from TCGA and checked for experimental design of Cancer vs. Normal for 19 kinds of epithelial cancers. We finally obtained 106 unique clusters for these cancer types. A cluster was designated to be homogeneous if at least half of its members show the same direction of expression pattern (either up- or down-regulated). For each cluster, we computed the homogeneous fraction as that of co-dysregulation throughout all cancer types analyzed. A significant P-value for this fraction was calculated as follow: for each cancer type, the expression of all its associated miRNAs was distributed randomly within these miRNAs for 10,000 times, keeping the distribution of up- and down-regulated miRNAs constant for each step. The homogeneous fraction over all cancers was computed, which yields the P-value as the number of sampled homogeneous fractions exceeding the original homogeneous fractions divided by 10,000.

In order to check whether clustered miRNAs are more enriched in cancer development compared to single miRNA, we calculated an enrichment score of log-odds (LOD) score for each cancer type:

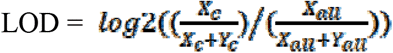

Where *X*_*c*_ and *Y*_*c*_ denote the number of cluster miRNAs and non-cluster miRNAs for each cancer type, and *X*_*all*_ denotes the number of the cluster miRNAs and denotes the number of the miRNAs not contained in any cluster. Here we take into account all known human miRNAs annotated in the human genome, thus designate as the 688 clustered miRNAs and 900 non-clustered miRNAs. In this case, a positive LOD score indicates enrichment for cluster miRNAs compared to non-cluster miRNAs in a specific cancer.

## Implementation and results

### Database content

In the current release of dbDEMC, the data of miRNA transcriptome of total 46,388 samples from 403 studies of human, mouse or rat were collected from public resources (Table S2). These profiles are derived from 149 subtypes of 40 different cancers and many different cell lines (Table S3). We then performed a systematic analysis on each dataset, and yielded a total of 807 experiments for differential expression analysis. dbDEMC 3.0 now hosts a total of 3,268 differentially expressed miRNAs, among them, 2,584 are specific to human. A total of 160,799 miRNA variations related to cancers were deposited in our database. The detailed information about number of miRNAs, cancer types, datasets, and experiments for different species was presented in **Table 1**.

**Figure 2A** depicts the number of DEMs for each type of human cancer. The breast cancer presents a large number of DEMs that 1,833 up-regulated and 1,988 down-regulated DEMs had been identified. The number of DEMs from mouse and rat can be found in Figure S1. Whereas Figure 2B demonstrates the number of DEMs validated by low-throughput methods across major cancers, the brain cancer, colorectal cancer and breast cancer are top ranked cancer types. Figure 2C shows the percentages of experiments for top ranked cancers. The breast cancer accounted 15% of the total experiments and ranked the first of the list. It was followed by colorectal cancer and lung cancer. Whereas the 9 different comparisons, cancer samples vs. normal controls constitutes about half of the total experiments, and followed by comparisons of high-grade versus low-grade cancer samples (Figure 2D). Overall, the sizes of analysis experiments and related literatures in dbDEMC 3.0 have a two-fold increment by comparing with the previous version (Figure S2).

**Figure 2.**
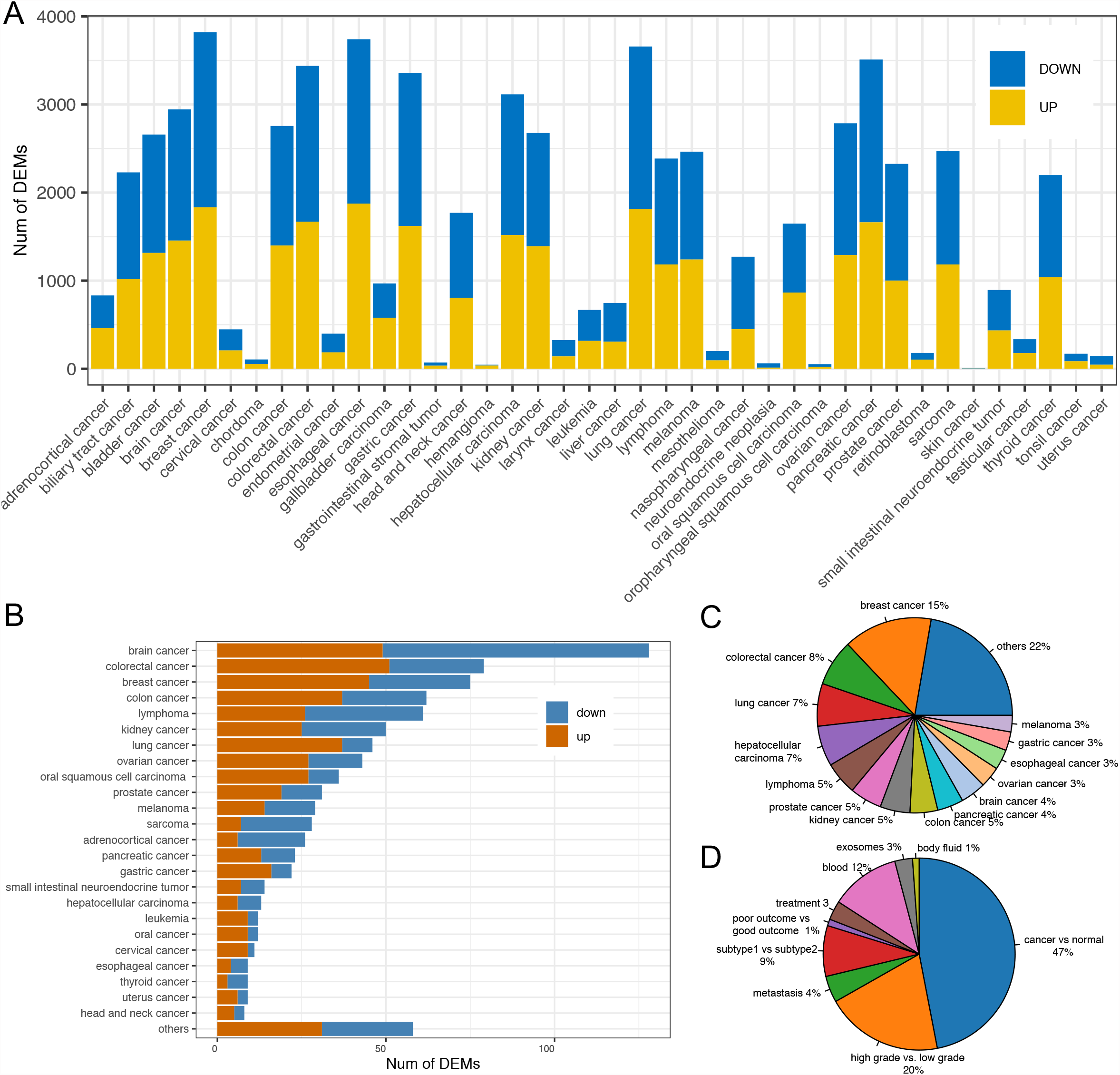
Statistics of data content in dbDEMC 3.0 for humans. **A**. Number of DEMs from each cancer types identified by high-throughput methods. **B**. Number of DEMs from major cancer types identified by low-throughput methods. **C**. The percentage of experiments for major cancer types. **D**. The percentage of experiments in seven types of experimental design.

### New features

In the dbDEMC 2.0, we assigned the different experimental designs to 7 different classes: Cancer vs. Normal; High grade vs. Low grade; Metastasis vs. Primary cancer; Subtype1 vs. Subtype2; Poor outcome vs. Good outcome; Blood sample of patients vs. Blood sample of normal controls, and also Treatment vs. Non-treatment. In recent years, many studies disclosed exosomes and microvesicles act as cell communication agents, where miRNA is the most important molecular in exosomes and microvesicles for regulating cancer progression [29]. In addition, circulating miRNAs were also widely found in body fluids and represent a gold mine of noninvasive biomarkers in cancer [30]. In this update version, we thus added these two classes of experimental design: Exosome sample from patients vs. Exosome sample from control, and Body fluid from patients vs. Body fluid from control. Moreover, for each DEMs set, targets of miRNAs and enrichment information of the target genes in the KEGG pathways and GO terms were deposited in the dbDEMC 3.0, which makes it possible for inspecting functional mechanisms behind a set of miRNAs.

### Newly designed web interface

The web interface of dbDEMC 3.0 has been significantly refined and improved, allowing better use of the data. The Search page permits users to perform a quick search and extract summarized information of a DEM list across cancer types. Users can also specify the cancer type, experimental design or platform to select the interested experiments (**Figure 3A–C**). After filtering the experimental results, users can select interested experiments. Detailed information of DEM related experiment, which includes description with the up-regulated and down-regulated miRNAs, can be accessed. In the functional chart section, heatmap of the DMEs expression, miRNA-target regulatory network for top ranked DEMs, as well as the bubble chart for miRNA targets enriched KEGG and GO terms were presented (Figure 3D). Using a single miRNA query, summary information of the interested miRNA can be retrieved, including the general description of the interested miRNA and differential expression summary heatmap which decided by number of experiments shows up expression or down expression. In addition, summary statistics tables for both high-throughput data analysis and low-throughput validation data were also displayed (Figure 3E).

**Figure 3.**
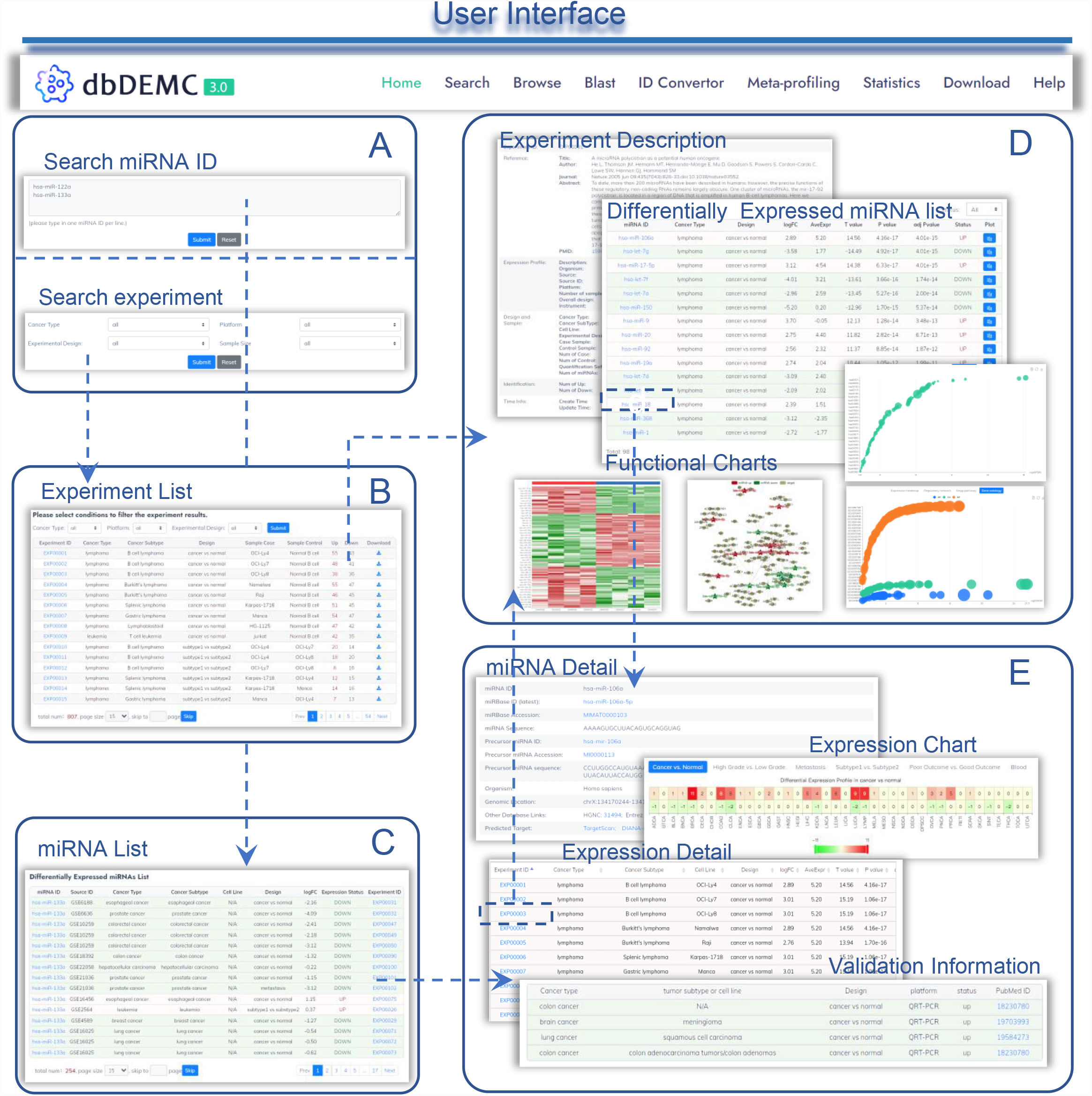
Web interface of dbDEMC 3.0. **A**. Search page. miRNAs can be searched via miRBase IDs, or filter experiments with interested conditions. **B**. Filtering result page of experiments. **C**. Search result page with example miRNAs. **D**. Experiment page. The page summarizes the description of the experiments and associated differentially expressed miRNA list, functional chart for expression heatmap, regularly network and miRNA targets enriched KEGG pathways and GO terms are also depicted. **E**. miRNA page. This page mainly consists of four sections: miRNA Summary, Expression Profile and Expression Detail and Validation.

### Analysis tools

miRBase is the central reference database for miRNA annotation by assigning names and unique gene IDs for novel miRNAs. During its development, some miRNA definition and annotation may have been changed. This leads to the inconsistence of the miRNA IDs from different data sets, which derived from different miRBase versions and making it difficult for comparing research results for integrative analysis. To solve this problem, we provide a “ID convertor” in our database, by which users could convert miRBase old version IDs to the latest version (v22.0) for the three species of human, mouse, and rat. In addition, other analyzing tools including BLAST and meta-profiling, which are used for sequence similarity search of unknown miRNAs and identify the confident cancer related miRNAs in pan-cancer wide, are also available in dbDEMC v3.0. For the meta-profiling study, the vote-counting approach was used to calculate the consistent score of differential expression for meta-analysis [31]. Common miRNAs identified in multiple cancer types with similar differential expression pattern may suggest they could have similar regulatory mechanisms and play important roles in cancer development.

### miRNA clusters are significantly overrepresented in cancers

A large proportion of miRNAs localized as conserved clusters in the genome and present similar expression pattern across tissues. It is critical to understand whether miRNA clusters present similar differential expression across cancers and correlate with the similar pathobiology. Here we obtained human miRNA cluster annotation from public resources, which includes 22.8% (688/2588) of mature miRNAs appear as 143 clusters of at least two members within (Table S4). We systematically analyzed the homogeneity of expression patterns within miRNA clusters. We excluded those clusters having less than half of all miRNAs annotated from the results of TCGA, which leads to 106 remaining clusters. The clusters are denoted as exhibiting a homogeneous expression pattern if annotated miRNA members are either up- or down-regulated (see Data processing). In total, cancer-associated clusters revealed homogeneous expression patterns for 74% of all annotated cancer, which confirms the hypothesis of co-regulation pattern of miRNA clusters in cancer. For example, the cluster of miR-142-5p, miR-142-3p, miR-4736 presents a consistent differential expression pattern in 91% (11/12) of the cancers analyzed (Table S5). A null model by randomly linking miRNA expression patterns (permutation 10,000 times within each cancer) indicated 52 clusters (49%, P-value < 0.05) showed a significantly higher homogeneity pattern in all 19 kinds of cancer compared to that expected by chance (Table S5). These clusters exhibit a homogeneous expression pattern in at least 78.5% of all these types of cancer.

To further investigate the association of miRNA clusters with different kinds of cancer, we estimated the enrichment of miRNA clusters in cancer-associated miRNAs by using a LOD score. We found enrichment for all 19 kinds of cancer (**Figure 4**). Within these 19 kinds of cancer, miRNAs located in clusters are, on average, 1.56 times (LOD = 0.65) enriched compared to random. In summary, our analyses show a significant enrichment of clustered miRNAs in cancers compare to the single miRNA members, which demonstrated that the different miRNAs within cluster act synergistically in cancer development.

**Figure 4.**
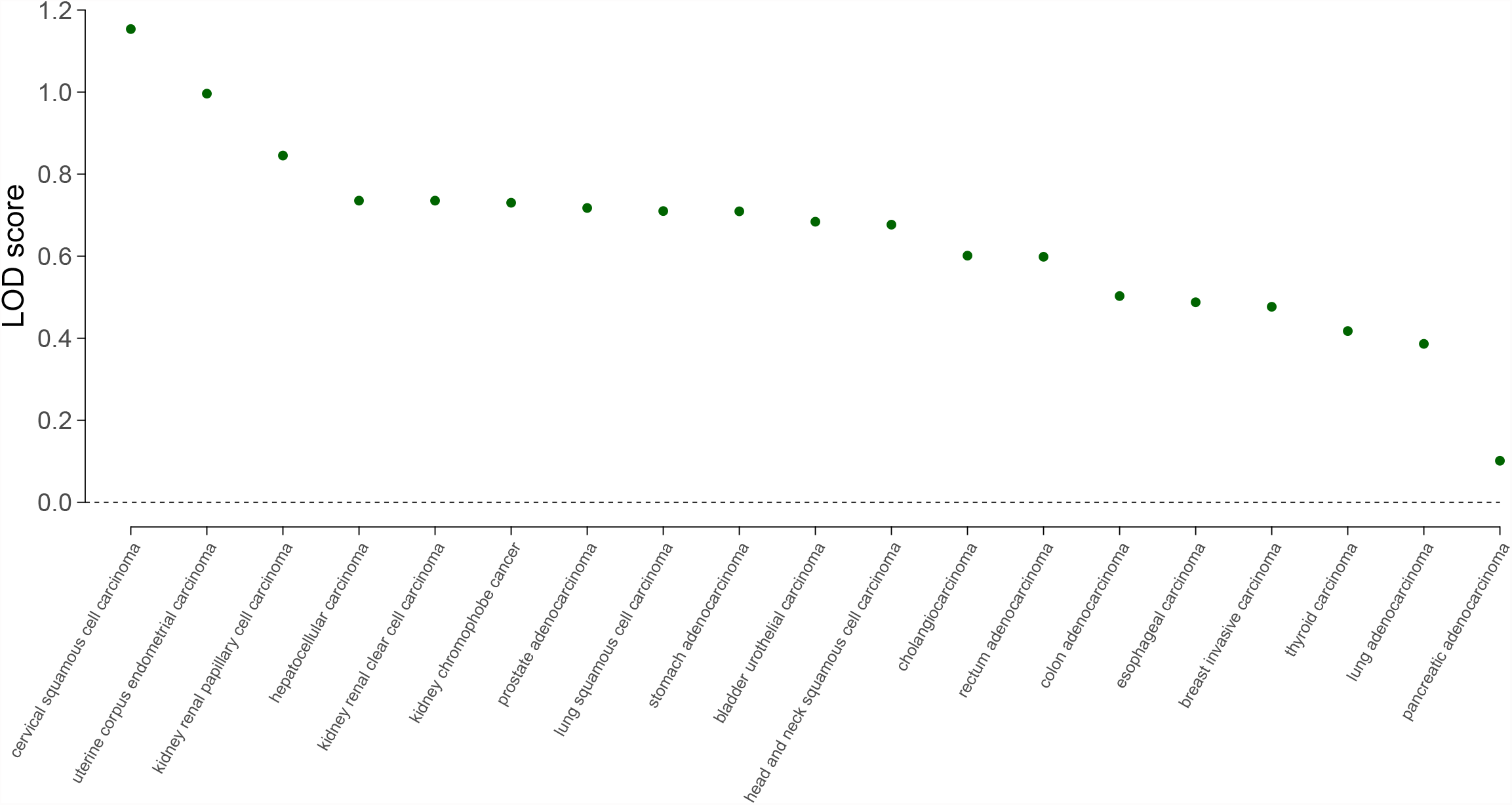
miRNA cluster enrichment for 19 kinds of cancer. For each cancer type, the log-odds (LOD) score is plotted. There is an enrichment of miRNA cluster members for all 19 kinds of cancer (100%, P-value < 1e-4). Within these 19 types of cancer, miRNAs located in clusters are, on average, 1.56 times (LOD = 0.65) enriched compared to random.

## Discussion

Over the last decade, a large amount of miRNA transcriptome profiles of various cancers have been generated. Many studies have performed miRNA transcriptome analysis to explore the underlying molecular mechanisms of miRNA genes in cancer development [32, 33]. This progress motivated a novel release of dbDEMC to keep track of the latest published data. Along with this, we curate these data and provide a platform to facilitate the study of miRNA-cancer associations. For dbDEMC 3.0, it not only contains more miRNA-cancer associations, we also extended our database to the species of mouse and rat, which benefits those studies characterizing the miRNA functional machinery in cancer using the model organisms. Beyond the rapid increase of data amount, our database now offers many new features and powerful tools for the downstream analysis of DEMs, such as the integrated target identification and functional enrichment analysis for miRNA-regulated biological processes.

One of the key questions of differential expression analysis of miRNAs is which cancer types are regulated by a particular miRNA (miRNA-centric view), or conversely, which miRNAs may involve in a given type of cancer (cancer-centric view). Our databases support both miRNA- and cancer-centric investigations, users are able to search miRNAs to determine the spectrum of cancer types that it involved, or to find candidate miRNA list based on reference of the miRBase which link to individual type of cancer. Although studies have indicated false positive and negative records may exist in miRBase, researchers need to be cautious about resources based on references from miRBase [34, 35]. Overall, we expect that dbDEMC 3.0 could serve as a valuable resource with comprehensive data amount and data analysis tools to facilitate the study of DEMs in cancers. In the future, we will continue to make improvements to the functional characterizations as well as integrate more heterogeneous data for the flexible analysis of miRNA functions in various cancers. We believe that the development of dbDEMC database can help accelerate the integration between miRNANome and cancer studies.

## Data availability

dbDEMC v3.0 is freely accessible at https://www.biosino.org/dbDEMC/.

## CRediT author statement

**Feng Xu:** Data curation, Formal analysis. **Yifan Wang:** Methodology, Validation. **Yunchao Ling:** Visualization. **Chenfen Zhou:** Data curation. **Haizhou Wang:** Data curation. **Andrew E. Teschendorff:** Methodology, Software. **Yi Zhao:** Investigation, Software. **Haitao Zhao:** Validation. **Yungang He:** Supervision, Investigation. **Guoqing Zhang:** Methodology, Supervision. **Zhen Yang:** Conceptualization, Supervision, Writing-Original draft. All authors read and approved the final manuscript.

## Acknowledgments

This work is supported by National Natural Science Foundation of China [91959106, 31871255, 91731310, and 81827901]; Strategic Priority Research Program of the Chinese Academy of Sciences [XDB38030100]; Shanghai Municipal Science and Technology [2017SHZDZX01].

## Tables

**Table 1 Summary of the data content of the current release of dbDEMC**

## Supplementary material

**Figure S1.**
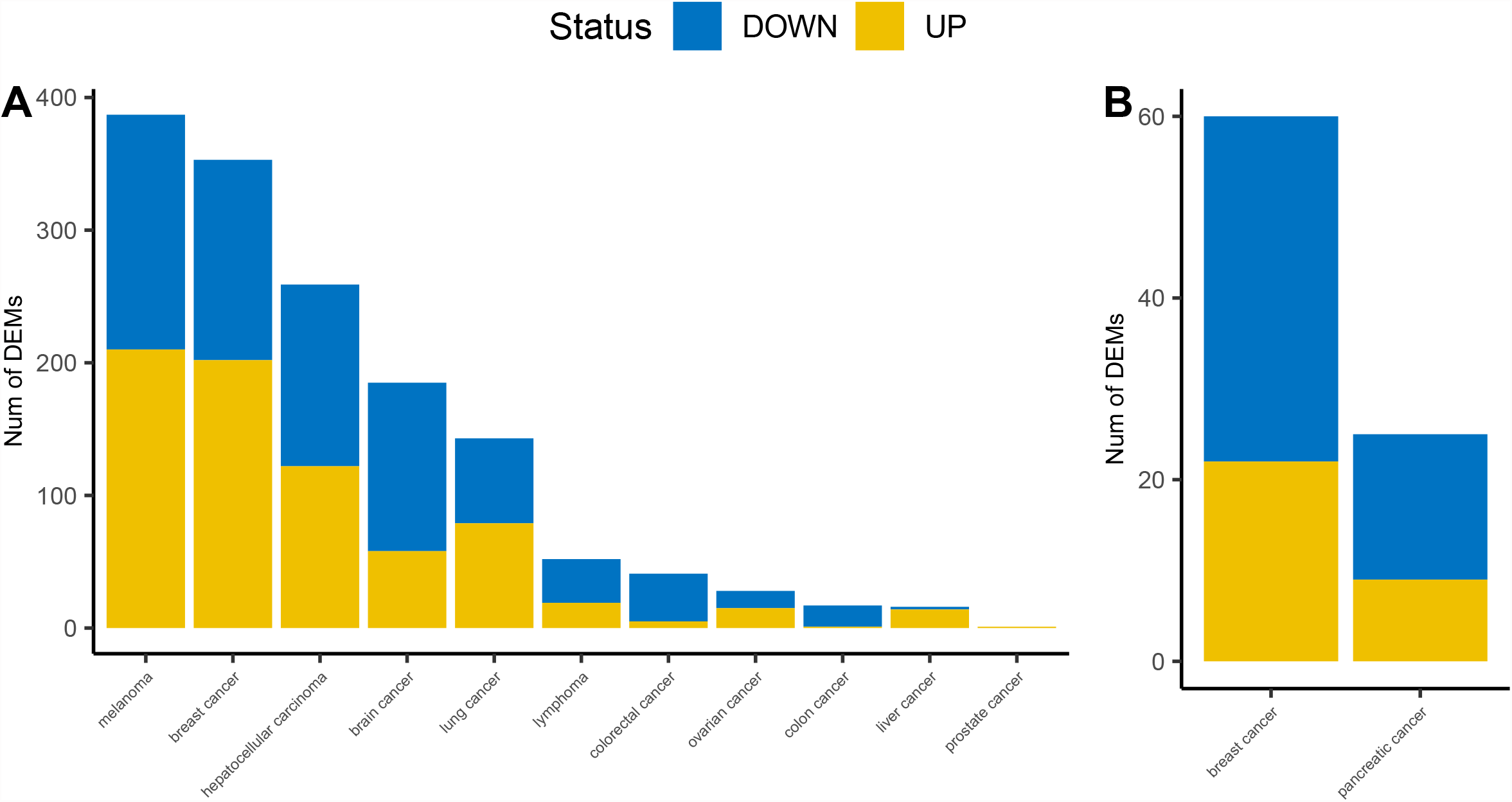
Number of differentially expressed miRNAs identified by high-throughput methods for each cancer type. **A**. Number of DEMs for mouse; **B**. Number of DEMs for rat.

**Figure S2.**
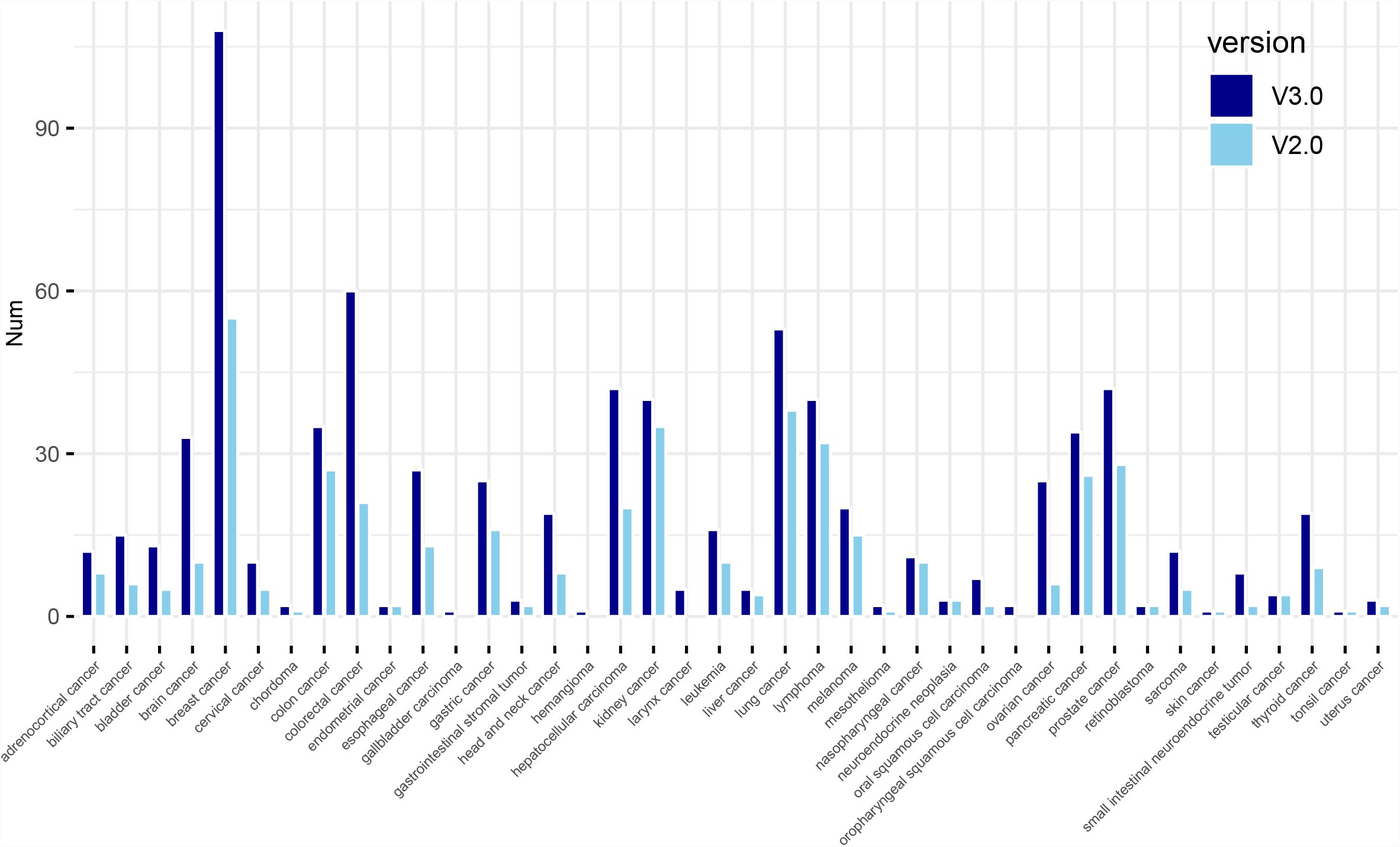
Increasing number of experiments for each cancer type. The number of experiments for each cancer type in dbDEMC v3.0 and v2.0 are depicted.

**Table S1 Adapters for miRNA-seq kits for the Illumina platform**.

**Table S2 The table below lists datasets collected from public resources, including the GEO, ArrayExpress, SRA and TCGA**. The Source Data ID, PubMed ID, Species, Cancer Type, platform ID and the total number of samples for each dataset were listed.

**Table S3 Cancer types and associated subtypes and cell line names covered in dbDEMC 3.0**.

**Table S4 All miRNA clusters and associated miRNA IDs identified**. A total of 143 miRNA clusters and associated 688 human mature miRNAs were obtained in the human genome.

**Table S5 miRNA clusters showing a homogenous expression pattern in all 19 kinds of cancer. T:** number of cancers for which each cluster showing a homogeneous expression pattern; **F:** number of diseases for which each cluster shows no homogeneous expression pattern; **Homogeneous-fraction:** number of diseases for which each cluster shows a homogeneous expression pattern for each miRNA cluster as defined; **P-value:** estimate by randomly linking miRNA-expression patterns 10,000 times within each cancer.

## References

[1] Ambros V. microRNAs: tiny regulators with great potential. Cell 2001;107:823–6.

[2] Bartel DP. MicroRNAs: genomics, biogenesis, mechanism, and function. Cell 2004;116:281–97.

[3] Kozomara A, Birgaoanu M, Griffiths-Jones S. miRBase: from microRNA sequences to function. Nucleic Acids Res 2019;47:D155–D62.

[4] Shivdasani RA. MicroRNAs: regulators of gene expression and cell differentiation. Blood 2006;108:3646–53.

[5] Hwang HW, Mendell JT. MicroRNAs in cell proliferation, cell death, and tumorigenesis. Br J Cancer 2007;96 Suppl:R40–4.

[6] Esquela-Kerscher A, Slack FJ. Oncomirs - microRNAs with a role in cancer. Nat Rev Cancer 2006;6:259–69.

[7] Wang J, Chen J, Sen S. MicroRNA as Biomarkers and Diagnostics. J Cell Physiol 2016;231:25–30.

[8] Wang H, Peng R, Wang J, Qin Z, Xue L. Circulating microRNAs as potential cancer biomarkers: the advantage and disadvantage. Clin Epigenetics 2018;10:59.

[9] Huang Z, Shi J, Gao Y, Cui C, Zhang S, Li J, et al. HMDD v3.0: a database for experimentally supported human microRNA-disease associations. Nucleic Acids Res 2019;47:D1013–D7.

[10] Xie B, Ding Q, Han H, Wu D. miRCancer: a microRNA-cancer association database constructed by text mining on literature. Bioinformatics 2013;29:638–44.

[11] Wang D, Gu J, Wang T, Ding Z. OncomiRDB: a database for the experimentally verified oncogenic and tumor-suppressive microRNAs. Bioinformatics 2014;30:2237–8.

[12] Yang Z, Ren F, Liu C, He S, Sun G, Gao Q, et al. dbDEMC: a database of differentially expressed miRNAs in human cancers. BMC Genomics 2010;11 Suppl 4:S5.

[13] Yang Z, Wu L, Wang A, Tang W, Zhao Y, Zhao H, et al. dbDEMC 2.0: updated database of differentially expressed miRNAs in human cancers. Nucleic Acids Res 2017;45:D812–D8.

[14] Barrett T, Wilhite SE, Ledoux P, Evangelista C, Kim IF, Tomashevsky M, et al. NCBI GEO: archive for functional genomics data sets--update. Nucleic Acids Res 2013;41:D991–5.

[15] Athar A, Fullgrabe A, George N, Iqbal H, Huerta L, Ali A, et al. ArrayExpress update - from bulk to single-cell expression data. Nucleic Acids Res 2019;47:D711–D5.

[16] Kodama Y, Shumway M, Leinonen R, International Nucleotide Sequence Database C. The Sequence Read Archive: explosive growth of sequencing data. Nucleic Acids Res 2012;40:D54–6.

[17] Zhao S, Gordon W, Du S, Zhang C, He W, Xi L, et al. QuickMIRSeq: a pipeline for quick and accurate quantification of both known miRNAs and isomiRs by jointly processing multiple samples from microRNA sequencing. BMC Bioinformatics 2017;18:180.

[18] Chen C, Khaleel SS, Huang H, Wu CH. Software for pre-processing Illumina next-generation sequencing short read sequences. Source Code Biol Med 2014;9:8.

[19] Zhong X, Heinicke F, Lie BA, Rayner S. Accurate Adapter Information Is Crucial for Reproducibility and Reusability in Small RNA Seq Studies. Noncoding RNA 2019;5.

[20] Langmead B, Trapnell C, Pop M, Salzberg SL. Ultrafast and memory-efficient alignment of short DNA sequences to the human genome. Genome Biol 2009;10:R25.

[21] Friedlander MR, Mackowiak SD, Li N, Chen W, Rajewsky N. miRDeep2 accurately identifies known and hundreds of novel microRNA genes in seven animal clades. Nucleic Acids Res 2012;40:37–52.

[22] Law CW, Chen Y, Shi W, Smyth GK. voom: Precision weights unlock linear model analysis tools for RNA-seq read counts. Genome Biol 2014;15:R29.

[23] Chu A, Robertson G, Brooks D, Mungall AJ, Birol I, Coope R, et al. Large-scale profiling of microRNAs for The Cancer Genome Atlas. Nucleic Acids Res 2016;44:e3.

[24] Ru Y, Kechris KJ, Tabakoff B, Hoffman P, Radcliffe RA, Bowler R, et al. The multiMiR R package and database: integration of microRNA-target interactions along with their disease and drug associations. Nucleic Acids Res 2014;42:e133.

[25] Karagkouni D, Paraskevopoulou MD, Chatzopoulos S, Vlachos IS, Tastsoglou S, Kanellos I, et al. DIANA-TarBase v8: a decade-long collection of experimentally supported miRNA-gene interactions. Nucleic Acids Res 2018;46:D239–D45.

[26] Huang HY, Lin YC, Li J, Huang KY, Shrestha S, Hong HC, et al. miRTarBase 2020: updates to the experimentally validated microRNA-target interaction database. Nucleic Acids Res 2020;48:D148–D54.

[27] Yu G, Wang LG, Han Y, He QY. clusterProfiler: an R package for comparing biological themes among gene clusters. OMICS 2012;16:284–7.

[28] Kabekkodu SP, Shukla V, Varghese VK, J Ds, Chakrabarty S, Satyamoorthy K. Clustered miRNAs and their role in biological functions and diseases. Biol Rev Camb Philos Soc 2018;93:1955–86.

[29] Sun Z, Shi K, Yang S, Liu J, Zhou Q, Wang G, et al. Effect of exosomal miRNA on cancer biology and clinical applications. Mol Cancer 2018;17:147.

[30] Cortez MA, Bueso-Ramos C, Ferdin J, Lopez-Berestein G, Sood AK, Calin GA. MicroRNAs in body fluids--the mix of hormones and biomarkers. Nat Rev Clin Oncol 2011;8:467–77.

[31] Guan P, Yin Z, Li X, Wu W, Zhou B. Meta-analysis of human lung cancer microRNA expression profiling studies comparing cancer tissues with normal tissues. J Exp Clin Cancer Res 2012;31:54.

[32] Ueda T, Volinia S, Okumura H, Shimizu M, Taccioli C, Rossi S, et al. Relation between microRNA expression and progression and prognosis of gastric cancer: a microRNA expression analysis. Lancet Oncol 2010;11:136–46.

[33] Iorio MV, Croce CM. MicroRNA dysregulation in cancer: diagnostics, monitoring and therapeutics. A comprehensive review. EMBO Mol Med 2012;4:143–59.

[34] Fromm B, Keller A, Yang X, Friedlander MR, Peterson KJ, Griffiths-Jones S. Quo vadis microRNAs? Trends Genet 2020;36:461–3.

[35] Alles J, Fehlmann T, Fischer U, Backes C, Galata V, Minet M, et al. An estimate of the total number of true human miRNAs. Nucleic Acids Res 2019;47:3353–64.

