## Supplementary material for "dbDEMC 3.0: Functional Exploration of Differentially Expressed miRNAs in Cancers of Human and Model Organisms": Table 1

**Table 1 Summary of the data content of the current release of dbDEMC**

|  | **miRNAs** | **Cancer Types** | **Cancer SubTypes** | **Datasets** | **Experiments** | **Samples** |
| --- | --- | --- | --- | --- | --- | --- |
| Homo sapiens | 2,584 | 40 | 149 | 373 | 763 | 45,974 |
| Mus musculus | 610 | 11 | 7 | 28 | 40 | 383 |
| Rattus norvegicus | 74 | 2 | 1 | 2 | 4 | 31 |
| All | 3,268 | 40 | 149 | 403 | 807 | 46,388 |
