## Supplementary material for "dbDEMC 3.0: Functional Exploration of Differentially Expressed miRNAs in Cancers of Human and Model Organisms": Table S1

**Table S1 Adapters for miRNA-seq kits for the Illumina platform.**

| Kit | Vendor | 3' Adapter sequence |
| --- | --- | --- |
| TruSeq Small RNA Library Preparation Kit | Illumina | 5'-TGGAATTCTCGGGTGCCAAGG-3' |
| NEXTflex Small RNA Sequencing Kit ** | PerkinElmer | 5'-TGGAATTCTCGGGTGCCAAGG-3' |
| NEBNext Multiplex Small RNA Library Prep Kit for Illumina | New England Biolabs | 5'-AGATCGGAAGAGCACACGTCT-3' |
| TailorMix miRNA Sample Preparation Kit | SeqMatic | 5'-TGGAATTCTCGGGTGCCAAGG-3' |
| CleanTag Small RNA Library Prep Kit | TriLink | 5'-TGGAATTCTCGGGTGCCAAGG-3' |
| QIAseq miRNA Library Kit | Qiagen | 5'-AACTGTAGGCACCATCAAT-3' |
| Small RNA-Seq Library Prep Kit | Lexogen | 5'-TGGAATTCTCGGGTGCCAAGGAACTCCAGTCAC-3' |
| SMARTer smRNA-Seq Kit for Illumina | Takara | 5'-AAAAAAAAAA-3' |
| CATS small RNA-seq Kit | Diagenode | 5'-GATCGGAAGAGCACACGTCTG-3' |
