## Supplementary material for "dbDEMC 3.0: Functional Exploration of Differentially Expressed miRNAs in Cancers of Human and Model Organisms": Table S2

**Table S2 The table below lists datasets collected from public resources, including the GEO, ArrayExpress, SRA and TCGA.** The Source Data ID, PubMed ID, Species, Cancer Type, platform ID and the total number of samples for each dataset were listed.

| Source Data ID | PubMed ID | Species | Cancer Type | GEO platform | Number Of Samples |
| --- | --- | --- | --- | --- | --- |
| GSE2399 | 15944707 | Homo sapiens | lymphoma, leukemia | GPL1899 | 40 |
| GSE2564 | 15944708 | Homo sapiens | breast cancer, renal carcinoma, lung cancer, prostate cancer, uterus carcinoma | GPL1986 | 334 |
| GSE4589 | 16784538 | Homo sapiens | breast cancer | GPL3238 | 20 |
| GSE5244 | 17243163 | Homo sapiens | uterus cancer | GPL3879 | 51 |
| GSE6188 | 18172293 | Homo sapiens | esophageal cancer | GPL4508 | 257 |
| GSE6636 |  | Homo sapiens | prostate cancer | GPL3238 | 47 |
| GSE7072 | 18026111 | Homo sapiens | retinoblastoma | GPL4879 | 6 |
| GSE7842 | 17922911 | Homo sapiens | breast cancer | GPL5173 | 119 |
| GSE8126 | 18676839 | Homo sapiens | prostate cancer | GPL5180 | 76 |
| GSE10259 | 19843336 | Homo sapiens | colorectal cancer | GPL4411 | 66 |
| GSE10694 | 18649363 | Homo sapiens | hepatocellular carcinoma | GPL6542 | 166 |
| GSE11016 | 18519660 | Homo sapiens | kidney cancer | GPL6668 | 32 |
| GSE11163 | 19351747 | Homo sapiens | head and neck cancer | GPL6690 | 27 |
| GSE12105 | 19228262 | Homo sapiens | kidney cancer | GPL6955 | 24 |
| GSE12303 | 18973228 | Homo sapiens | brain cancer | GPL7114 | 48 |
| GSE12933 | 19047678 | Homo sapiens | lymphomas | GPL7373 | 8 |
| GSE13030 |  | Homo sapiens | brain cancer | GPL7389 | 93 |
| GSE14857 | 19676045 | Homo sapiens | prostate cancer | GPL6955 | 24 |
| GSE15008 |  | Homo sapiens | lung cancer | GPL8176 | 375 |
| GSE16025 | 19584273 | Homo sapiens | lung cancer | GPL5106 | 71 |
| GSE16456 | 19737949 | Homo sapiens | esophageal cancer | GPL6955 | 32 |
| GSE16558 |  | Homo sapiens | lymphoma | GPL8695 | 65 |
| GSE17792 | 19703993 | Homo sapiens | brain cancer | GPL7436 | 17 |
| GSE20099 | 21128279 | Homo sapiens | esophageal cancer | GPL8871 | 51 |
| GSE6857 | 18176954 | Homo sapiens | hepatocellular carcinoma | GPL4700 | 482 |
| GSE7055 | 18459106 | Homo sapiens | prostate cancer | GPL4700 | 114 |
| GSE7828 | 18230780 | Homo sapiens | colon cancer | GPL4700 | 170 |
| GSE15885 | 19432961 | Homo sapiens | breast cancer | GPL6127 | 29 |
| GSE17498 | 19846888 | Homo sapiens | lymphoma | GPL8227 | 102 |
| GSE18509 | 20304954 | Homo sapiens | melanoma | GPL9081 | 16 |
| GSE18392 | 19922656 | Homo sapiens | colon cancer | GPL8178 | 145 |
| GSE19856 | 20484036 | Homo sapiens | adrenocortical cancer | GPL8227 | 30 |
| GSE19387 | 20357817 | Homo sapiens | melanoma | GPL9081 | 40 |
| GSE22058 | 20739924 | Homo sapiens | hepatocellular carcinoma | GPL10457 | 397 |
| GSE21036 | 20579941 | Homo sapiens | prostate cancer | GPL8227 | 142 |
| GSE23022 | 21400514 | Homo sapiens | prostate cancer | GPL8786 | 40 |
| GSE24390 | 21156648 | Homo sapiens | brain cancer | GPL7436 | 24 |
| GSE24485 | 21460242 | Homo sapiens | lymphoma | GPL8227 | 22 |
| GSE18546 |  | Homo sapiens | sarcoma | GPL7738 | 20 |
| GSE24996 | 21893020 | Homo sapiens | melanoma | GPL6955 | 31 |
| GSE19783 | 21364938 | Homo sapiens | breast cancer | GPL8227 | 216 |
| GSE20077 | 22056881 | Homo sapiens | hepatocellular carcinoma | GPL8227 | 11 |
| GSE19980 |  | Homo sapiens | hepatocellular carcinoma | GPL9954 | 12 |
| GSE21362 | 21298008 | Homo sapiens | hepatocellular carcinoma | GPL10312 | 146 |
| GSE28700 | 21703006 | Homo sapiens | gastric cancer | GPL9081 | 44 |
| GSE29352 |  | Homo sapiens | pancreatic cancer | GPL13606 | 43 |
| GSE26245 |  | Homo sapiens | prostate cancer | GPL11350 | 96 |
| GSE22816 | 21859927 | Homo sapiens | adrenocortical cancer | GPL13347 | 52 |
| GSE22216 | 21737487 | Homo sapiens | breast cancer | GPL8178 | 210 |
| GSE22420 | 21701491 | Homo sapiens | lymphoma | GPL10554 | 150 |
| GSE24508 | 21375733 | Homo sapiens | breast cancer | GPL10349 | 19 |
| GSE23383 | 21811625 | Homo sapiens | ovarian cancer | GPL9735 | 6 |
| GSE23739 | 21415212 | Homo sapiens | gastric cancer | GPL7731 | 80 |
| GSE32232 | 22110251 | Homo sapiens | lymphoma | GPL9081 | 46 |
| GSE25820 | 21953293 | Homo sapiens | pancreatic cancer | GPL7731 | 18 |
| GSE32678 | 22261810 | Homo sapiens | pancreatic cancer | GPL7723 | 32 |
| GSE25653 | 21435193 | Homo sapiens | melanoma | GPL6955 | 56 |
| GSE31741 | 22258453 | Homo sapiens | gastrointestinal stromal tumor | GPL10406 | 32 |
| GSE30454 | 21844009 | Homo sapiens | colorectal cancer | GPL8179 | 74 |
| GSE31377 | 21921041 | Homo sapiens | lymphoma | GPL8227 | 56 |
| GSE33961 |  | Homo sapiens | colorectal cancer | GPL7731 | 12 |
| GSE26323 | 23478189 | Homo sapiens | hepatocellular carcinoma | GPL11362 | 6 |
| GSE35579 |  | Homo sapiens | melanoma | GPL15183 | 71 |
| GSE31629 | 22264793 | Homo sapiens | leukemia | GPL7731 | 62 |
| GSE32906 |  | Homo sapiens | nasopharyngeal carcinoma | GPL11350 | 22 |
| GSE35412 | 22438871 | Homo sapiens | breast cancer | GPL9731 | 75 |
| GSE35602 | 22452939 | Homo sapiens | colorectal cancer | GPL8227 | 59 |
| GSE29965 | 22504665 | Homo sapiens | esophageal cancer | GPL7722 | 12 |
| GSE31045 | 22350414 | Homo sapiens | sarcoma | GPL7723 | 25 |
| GSE29135 | 22331473 | Homo sapiens | lung cancer | GPL8179 | 387 |
| GSE29491 | 21478429 | Homo sapiens | lymphoma | GPL8617 | 26 |
| GSE28955 |  | Homo sapiens | pancreatic cancer | GPL6955 | 23 |
| GSE37407 | 22294488 | Homo sapiens | breast cancer | GPL13703 | 61 |
| GSE28090 | 22102710 | Homo sapiens | lymphoma | GPL11487 | 25 |
| GSE36682 |  | Homo sapiens | nasopharyngeal carcinoma | GPL15311 | 68 |
| GSE35794 |  | Homo sapiens | endometrial cancer | GPL10850 | 22 |
| GSE30656 | 22330141 | Homo sapiens | cervical cancer | GPL6955 | 47 |
| GSE32960 | 23408429 | Homo sapiens | nasopharyngeal carcinoma | GPL14722 | 330 |
| GSE25631 | 22570426 | Homo sapiens | brain cancer | GPL8179 | 87 |
| GSE33743 | 22450781 | Homo sapiens | gastric cancer | GPL14895 | 41 |
| GSE36681 | 22573352 | Homo sapiens | lung cancer | GPL8179 | 206 |
| GSE28100 | 22761427 | Homo sapiens | oral squamous cell carcinoma | GPL10850 | 20 |
| GSE29248 | 22046296 | Homo sapiens | lung cancer | GPL8179 | 12 |
| GSE32957 | 22707408 | Homo sapiens | liver cancer | GPL14732 | 35 |
| GSE37527 | 22964023 | Homo sapiens | breast cancer | GPL14767 | 12 |
| GSE39678 | 23499894 | Homo sapiens | hepatocellular carcinoma | GPL15852 | 24 |
| GSE38389 | 22850566 | Homo sapiens | colorectal cancer | GPL11039 | 140 |
| GSE41282 | 23799849 | Homo sapiens | kidney cancer | GPL8786 | 38 |
| GSE28423 | 23133552 | Homo sapiens | sarcoma | GPL8227 | 23 |
| GSE40525 | 23125021 | Homo sapiens | breast cancer | GPL8227 | 120 |
| GSE34460 | 23111773 | Homo sapiens | melanoma | GPL15019 | 21 |
| GSE38781 | 23155457 | Homo sapiens | pancreatic cancer | GPL14903 | 19 |
| GSE33125 | 23028787 | Homo sapiens | colon cancer | GPL8179 | 18 |
| GSE33332 | 23239665 | Homo sapiens | kidney cancer | GPL10850 | 12 |
| GSE39486 | 22921398 | Homo sapiens | brain cancer | GPL15829 | 6 |
| GSE33232 |  | Homo sapiens | head and neck cancer | GPL8786 | 284 |
| GSE35982 | 22703586 | Homo sapiens | colorectal cancer | GPL14767 | 32 |
| GSE19945 |  | Homo sapiens | lung cancer | GPL9948 | 63 |
| GSE36802 | 23233736 | Homo sapiens | prostate cancer | GPL8786 | 42 |
| GSE40744 | 23390000 | Homo sapiens | hepatocellular carcinoma | GPL14613 | 76 |
| GSE33568 | 23328977 | Homo sapiens | small intestinal neuroendocrine tumor | GPL8786 | 25 |
| GSE45264 |  | Homo sapiens | lymphoma | GPL16770 | 22 |
| GSE45604 | 24518785 | Homo sapiens | prostate cancer | GPL14613 | 60 |
| GSE36915 | 22613005 | Homo sapiens | hepatocellular carcinoma | GPL8179 | 89 |
| GSE31164 | 23496901 | Homo sapiens | hepatocellular carcinoma | GPL10850 | 110 |
| GSE37053 | 23613318 | Homo sapiens | lymphoma | GPL8227 | 57 |
| GSE47764 | 24169343 | Homo sapiens | liver cancer | GPL11487 | 6 |
| GSE29742 | 23660872 | Homo sapiens | neuroendocrine neoplasia | GPL8227 | 54 |
| GSE26595 | 24222951 | Homo sapiens | gastric cancer | GPL8179 | 68 |
| GSE40355 |  | Homo sapiens | bladder cancer | GPL8227 | 48 |
| GSE47582 | 24189146 | Homo sapiens | kidney cancer | GPL8786 | 26 |
| GSE40345 | 23828229 | Homo sapiens | mesothelioma | GPL8179 | 31 |
| GSE41874 |  | Homo sapiens | hepatocellular carcinoma | GPL7722 | 10 |
| GSE45666 | 23722663 | Homo sapiens | breast cancer | GPL14767 | 116 |
| GSE49246 | 24239208 | Homo sapiens | colon cancer | GPL17496 | 80 |
| GSE32922 |  | Homo sapiens | breast cancer | GPL7723 | 38 |
| GSE41369 | 24120476 | Homo sapiens | pancreatic cancer | GPL16142 | 18 |
| GSE43796 | 24072181 | Homo sapiens | pancreatic cancer | GPL15159 | 31 |
| GSE29622 | 22362069 | Homo sapiens | colon cancer | GPL11162 | 65 |
| GSE35834 | 23987127 | Homo sapiens | colon cancer | GPL8786 | 158 |
| GSE42906 | 24121164 | Homo sapiens | lymphoma | GPL9081 | 55 |
| GSE44124 | 24098452 | Homo sapiens | breast cancer | GPL14767 | 53 |
| GSE47841 | 24512620 | Homo sapiens | ovarian cancer | GPL14613 | 30 |
| GSE50505 | 24298054 | Homo sapiens | kidney cancer | GPL17667 | 28 |
| GSE51908 | 25368993 | Homo sapiens | leukemia | GPL8786 | 190 |
| GSE54088 | 26069251 | Homo sapiens | colorectal cancer | GPL8178 | 34 |
| GSE54397 | 25167801 | Homo sapiens | gastric cancer | GPL15159 | 32 |
| GSE54492 | 25650662 | Homo sapiens | melanoma | GPL10262 | 25 |
| GSE31277 | 24209638 | Homo sapiens | head and neck cancer | GPL9770 | 42 |
| GSE38167 | 24479446 | Homo sapiens | breast cancer | GPL14943 | 67 |
| GSE40807 | 27036030 | Homo sapiens | thyroid cancer | GPL8227 | 80 |
| GSE41012 |  | Homo sapiens | colorectal cancer | GPL7724 | 35 |
| GSE41032 | 24519663 | Homo sapiens | brain cancer | GPL15516 | 6 |
| GSE41655 |  | Homo sapiens | colorectal cancer | GPL11487 | 107 |
| GSE43039 | 24606633 | Homo sapiens | nasopharyngeal carcinoma | GPL16414 | 40 |
| GSE44899 | 24917463 | Homo sapiens | breast cancer | GPL7723 | 80 |
| GSE45238 | 25351956 | Homo sapiens | oral squamous cell carcinoma | GPL8179 | 80 |
| GSE45364 | 24375455 | Homo sapiens | sarcoma | GPL16851 | 115 |
| GSE48088 | 25047087 | Homo sapiens | breast cancer | GPL14613 | 36 |
| GSE48267 | 24865442 | Homo sapiens | colon cancer | GPL10850 | 122 |
| GSE51853 | 24903339 | Homo sapiens | lung cancer | GPL7341 | 131 |
| GSE53870 |  | Homo sapiens | liver cancer | GPL18118 | 72 |
| GSE53992 | 25017828 | Homo sapiens | liver cancer | GPL18159 | 46 |
| GSE56183 |  | Homo sapiens | chordoma | GPL14613 | 12 |
| GSE56350 | 24921248 | Homo sapiens | colorectal cancer | GPL16744 | 104 |
| GSE57370 | 25670083 | Homo sapiens | kidney cancer | GPL16770 | 66 |
| GSE57555 | 26538415 | Homo sapiens | hepatocellular carcinoma | GPL18044 | 64 |
| GSE57768 | 25764000 | Homo sapiens | skin cancer | GPL17537 | 48 |
| GSE60978 | 25579086 | Homo sapiens | pancreatic cancer | GPL15159 | 89 |
| GSE61438 |  | Homo sapiens | breast cancer | GPL8179 | 69 |
| GSE62054 |  | Homo sapiens | thyroid cancer | GPL8179 | 25 |
| GSE63805 | 26134223 | Homo sapiens | lung cancer | GPL18410 | 62 |
| GSE64318 | 26089375 | Homo sapiens | prostate cancer | GPL8227 | 54 |
| GSE65819 | 26017449 | Homo sapiens | ovarian cancer | GPL19765 | 121 |
| GSE66274 | 26409826 | Homo sapiens | esophageal cancer | GPL19823 | 60 |
| GSE67354 |  | Homo sapiens | gastric cancer | GPL19952 | 10 |
| GSE69580 |  | Homo sapiens | hepatocellular carcinoma | GPL10850 | 10 |
| GSE70574 |  | Homo sapiens | colorectal cancer | GPL15018 | 16 |
| GSE73487 |  | Homo sapiens | colon cancer | GPL8786 | 113 |
| GSE74190 |  | Homo sapiens | lung cancer | GPL19622 | 136 |
| GSE75630 | 26867589 | Homo sapiens | tonsil cancer | GPL21198 | 46 |
| GSE76260 |  | Homo sapiens | prostate cancer | GPL8179 | 64 |
| GSE77380 |  | Homo sapiens | lung cancer, colorectal cancer, gastric cancer | GPL16770 | 108 |
| GSE80038 |  | Homo sapiens | breast cancer | GPL18044 | 20 |
| GSE24709 |  | Homo sapiens | lung cancer | GPL9040 | 71 |
| GSE25609 |  | Homo sapiens | colon cancer | GPL8179 | 77 |
| GSE27486 | 23029380 | Homo sapiens | lung cancer | GPL11432 | 45 |
| GSE31309 | 22242178 | Homo sapiens | breast cancer | GPL14132 | 105 |
| GSE31568 | 21892151 | Homo sapiens | pancreatic cancer, melanoma, prostate cancer, ovarian cancer, gastric cancer | GPL9040 | 453 |
| GSE31801 | 22235027 | Homo sapiens | ovarian cancer | GPL8179 | 121 |
| GSE38419 | 22871070 | Homo sapiens | kidney cancer | GPL9040 | 42 |
| GSE39845 | 23758639 | Homo sapiens | colon cancer | GPL14613 | 26 |
| GSE41321 | 23400111 | Homo sapiens | retinoblastoma | GPL148767 | 6 |
| GSE41922 | 23797906 | Homo sapiens | breast cancer | GPL16224 | 54 |
| GSE48137 |  | Homo sapiens | kidney cancer | GPL16770 | 102 |
| GSE55993 | 25344866 | Homo sapiens | lung cancer | GPL16770 | 80 |
| GSE59856 | 25706130 | Homo sapiens | pancreatic cancer, biliary tract cancer, colon cancer, gastric cancer, esophagus cancer, liver cancer | GPL18941 | 571 |
| GSE65071 | 25784290 | Homo sapiens | sarcoma | GPL19631 | 35 |
| GSE71043 |  | Homo sapiens | esophageal cancer | GPL18042 | 6 |
| GSE37406 |  | Homo sapiens | pancreatic cancer | GPL14903 | 24 |
| GSE46188 | 23635652 | Homo sapiens | pancreatic adenocarcinoma | GPL8179 | 42 |
| GSE50224 |  | Homo sapiens | esophageal cancer | GPL8786 | 36 |
| GSE53850 |  | Homo sapiens | lymphoma | GPL16770 | 8 |
| GSE62137 |  | Homo sapiens | chronic lymphocytic leukemia | GPL14767 | 44 |
| GSE63159 |  | Homo sapiens | gastrointestinal stromal tumor | GPL10656 | 34 |
| GSE67257 | 26252371 | Homo sapiens | liver cancer | GPL18941 | 20 |
| GSE74562 |  | Homo sapiens | pancreatic adenocarcinoma | GPL14613 | 8 |
| TCGA_BRCA |  | Homo sapiens | breast cancer | miRNA-seq | 1189 |
| TCGA_KIRP |  | Homo sapiens | kidney cancer | miRNA-seq | 324 |
| TCGA_KIRC |  | Homo sapiens | kidney cancer | miRNA-seq | 583 |
| TCGA_KICH |  | Homo sapiens | kidney cancer | miRNA-seq | 91 |
| TCGA_LUAD |  | Homo sapiens | lung cancer | miRNA-seq | 555 |
| TCGA_LUSC |  | Homo sapiens | lung cancer | miRNA-seq | 518 |
| TCGA_UCEC |  | Homo sapiens | uterus cancer | miRNA-seq | 621 |
| TCGA_COAD |  | Homo sapiens | colon cancer | miRNA-seq | 450 |
| TCGA_READ |  | Homo sapiens | colorectal cancer | miRNA-seq | 161 |
| TCGA_HNSC |  | Homo sapiens | head and neck cancer | miRNA-seq | 564 |
| TCGA_THCA |  | Homo sapiens | thyroid cancer | miRNA-seq | 566 |
| TCGA_PRAD |  | Homo sapiens | prostate cancer | miRNA-seq | 532 |
| TCGA_STAD |  | Homo sapiens | gastric cancer | miRNA-seq | 475 |
| TCGA_LIHC |  | Homo sapiens | liver cancer | miRNA-seq | 420 |
| TCGA_CESC |  | Homo sapiens | cervical cancer | miRNA-seq | 304 |
| TCGA_ACC |  | Homo sapiens | adrenocortical cancer | miRNA-seq | 80 |
| TCGA_PAAD |  | Homo sapiens | pancreatic cancer | miRNA-seq | 183 |
| TCGA_ESCA |  | Homo sapiens | esophageal cancer | miRNA-seq | 187 |
| TCGA_CHOL |  | Homo sapiens | liver cancer | miRNA-seq | 45 |
| TCGA_SKCM |  | Homo sapiens | skin cancer | miRNA-seq | 449 |
| TCGA_BLCA |  | Homo sapiens | bladder cancer | miRNA-seq | 426 |
| TCGA_TGCT |  | Homo sapiens | testicular cancer | miRNA-seq | 149 |
| GSE59247 | 27396337 | Homo sapiens | breast cancer | GPL15019 | 48 |
| GSE70534 | 27353039 | Homo sapiens | small intestinal neuroendocrine tumor | GPL17537 | 79 |
| GSE71905 | 26398221 | Homo sapiens | liver cancer | GPL15446 | 8 |
| GSE78775 | 27518872 | Homo sapiens | gastric cancer | GPL10850 | 56 |
| GSE75283 | 27775819 | Homo sapiens | hepatocellular carcinoma | GPL16298 | 65 |
| GSE70080 |  | Homo sapiens | lung cancer | GPL20591 | 54 |
| GSE83924 | 28289479 | Homo sapiens | colorectal cancer | GPL16384 | 40 |
| GSE51945 | 25952770 | Homo sapiens | lung cancer | GPL8786 | 6 |
| GSE51946 | 25952770 | Homo sapiens | lung cancer | GPL14613 | 18 |
| GSE63319 |  | Homo sapiens | brain cancer | GPL16384 | 18 |
| GSE73182 | 26871295 | Homo sapiens | thyroid cancer | GPL20194 | 24 |
| GSE74618 | 27614046 | Homo sapiens | hepatocellular carcinoma | GPL14613 | 240 |
| GSE39567 |  | Homo sapiens | breast cancer | GPL8179 | 56 |
| GSE53592 | 24498407 | Homo sapiens | colon cancer | GPL8786 | 6 |
| GSE78037 | 29100302 | Homo sapiens | leukemia | GPL7731 | 6 |
| GSE82064 |  | Homo sapiens | oropharyngeal squamous cell carcinoma | GPL21968 | 96 |
| GSE87715 | 28052011 | Homo sapiens | lymphoma | GPL20712 | 8 |
| GSE93415 | 28641313 | Homo sapiens | gastric cancer | GPL19071 | 40 |
| GSE69126 | 27816053 | Homo sapiens | larynx cancer | GPL15018 | 8 |
| GSE69127 | 27816053 | Homo sapiens | larynx cancer | GPL20228 | 16 |
| GSE94536 | 28514730 | Homo sapiens | lung cancer | GPL21536 | 9 |
| GSE71130 | 26325386 | Homo sapiens | colorectal cancer | GPL18509 | 35 |
| GSE87828 | 27916857 | Homo sapiens | lymphoma | GPL16384 | 95 |
| GSE95385 | 24703100 | Homo sapiens | kidney cancer | GPL16770 | 8 |
| GSE55625 |  | Homo sapiens | sarcoma | GPL18116 | 17 |
| GSE92595 | 28098823 | Homo sapiens | head and neck cancer | GPL20968 | 40 |
| GSE60117 | 27384993 | Homo sapiens | prostate cancer | GPL13264 | 75 |
| GSE97049 |  | Homo sapiens | esophageal cancer | GPL21572 | 14 |
| GSE81873 | 28716985 | Homo sapiens | ovarian cancer | GPL17485 | 32 |
| GSE98406 |  | Homo sapiens | colon cancer | GPL16384 | 35 |
| GSE86995 |  | Homo sapiens | breast cancer | GPL20906 | 68 |
| GSE99415 | 28819095 | Homo sapiens | gastric cancer | GPL18058 | 12 |
| GSE81350 | 28052030 | Homo sapiens | esophageal cancer | GPL21263 | 16 |
| GSE83693 | 27655640 | Homo sapiens | ovarian cancer | GPL22079 | 20 |
| GSE81581 | 27662660 | Homo sapiens | colorectal cancer | GPL16384 | 51 |
| GSE83270 | 28359916 | Homo sapiens | breast cancer | GPL22003 | 12 |
| GSE71533 | 26590090 | Homo sapiens | pancreatic cancer | GPL18058 | 52 |
| GSE101976 | 29042553 | Homo sapiens | ovarian cancer | GPL17904 | 95 |
| GSE99362 | 28716051 | Homo sapiens | mesothelioma | GPL8179 | 58 |
| GSE90603 | 30176243 | Homo sapiens | brain cancer | GPL21572 | 24 |
| GSE104165 | 30886199 | Homo sapiens | gallbladder carcinoma | GPL18402 | 48 |
| GSE93850 | 28931080 | Homo sapiens | brain cancer | GPL22948 | 30 |
| GSE106744 | 27916857 | Homo sapiens | leukemia | GPL16384 | 34 |
| GSE98463 | 29372649 | Homo sapiens | oral squamous cell carcinoma | GPL21572 | 16 |
| GSE108153 | 30588338 | Homo sapiens | colorectal cancer | GPL13264 | 42 |
| GSE57897 | 31209222 | Homo sapiens | breast cancer | GPL18722 | 453 |
| GSE97070 | 29298817 | Homo sapiens | thyroid cancer | GPL18402 | 20 |
| GSE112009 |  | Homo sapiens | brain cancer | GPL21572 | 15 |
| GSE112743 |  | Homo sapiens | cervical cancer | GPL18402 | 6 |
| GSE113749 |  | Homo sapiens | lymphoma | GPL21942 | 95 |
| GSE113956 | 30867806 | Homo sapiens | oral squamous cell carcinoma | GPL18058 | 40 |
| GSE115016 | 30364632 | Homo sapiens | hepatocellular carcinoma | GPL21572 | 24 |
| GSE115513 | 26740022 | Homo sapiens | colorectal cancer | GPL18402 | 1399 |
| GSE109421 | 29894486 | Homo sapiens | lymphoma | GPL8227 | 29 |
| GSE81137 |  | Homo sapiens | cervical cancer | GPL16384 | 12 |
| GSE110402 | 29902577 | Homo sapiens | colorectal cancer | GPL14613 | 81 |
| GSE86277 | 30115973 | Homo sapiens | breast cancer | GPL14613 | 71 |
| GSE86278 | 30115973 | Homo sapiens | breast cancer | GPL16384 | 45 |
| GSE72281 |  | Homo sapiens | colorectal cancer | GPL18058 | 12 |
| GSE103317 |  | Homo sapiens | small intestinal neuroendocrine tumor | GPL10406 | 54 |
| GSE116251 | 30201497 | Homo sapiens | kidney cancer | GPL25243 | 36 |
| GSE106817 | 30333487 | Homo sapiens | breast cancer, colorectal cancer, esophageal cancer, gastric cancer, hepatocellular carcinoma, lung cancer, sarcoma, ovarian cancer, pancreatic cancer | GPL21263 | 4046 |
| GSE113486 | 30382619 | Homo sapiens | breast cancer, colorectal cancer, esophageal cancer, gastric cancer, hepatocellular carcinoma, lung cancer, sarcoma, ovarian cancer, pancreatic cancer, brain cancer, prostate cancer, bladder cancer, biliary tract cancer | GPL21263 | 972 |
| GSE119268 | 30404194 | Homo sapiens | lung cancer | GPL18402 | 155 |
| GSE113234 | 30452625 | Homo sapiens | prostate cancer | GPL19730 | 87 |
| GSE126093 | 31300733 | Homo sapiens | colorectal cancer | GPL18058 | 20 |
| GSE112840 | 30719423 | Homo sapiens | esophageal cancer | GPL23365 | 104 |
| GSE115108 | 30737378 | Homo sapiens | colon cancer | GPL18058 | 9 |
| GSE113629 | 30799952 | Homo sapiens | thyroid cancer | GPL24741 | 10 |
| GSE128446 |  | Homo sapiens | colorectal cancer | GPL14767 | 22 |
| GSE119055 | 30823895 | Homo sapiens | ovarian cancer | GPL21572 | 9 |
| GSE109943 | 30951707 | Homo sapiens | chordoma | GPL21572 | 16 |
| GSE132619 | 31242597 | Homo sapiens | colorectal cancer | GPL21572 | 8 |
| GSE118613 | 31159814 | Homo sapiens | nasopharyngeal cancer | GPL19888 | 150 |
| GSE117666 | 31227006 | Homo sapiens | melanoma | GPL16384 | 6 |
| GSE120930 | 31269432 | Homo sapiens | breast cancer | GPL15018 | 9 |
| GSE124678 |  | Homo sapiens | larynx cancer | GPL16770 | 37 |
| GSE118782 |  | Homo sapiens | breast cancer | GPL8786 | 40 |
| GSE137865 |  | Homo sapiens | oral squamous cell carcinoma | GPL8786 | 18 |
| GSE104717 |  | Homo sapiens | bladder cancer | GPL21572 | 6 |
| GSE138764 | 29221210 | Homo sapiens | brain cancer | GPL18402 | 42 |
| GSE108724 | 31485608 | Homo sapiens | hepatocellular carcinoma | GPL20712 | 14 |
| GSE135518 | 31533233 | Homo sapiens | sarcoma | GPL20712 | 64 |
| GSE142699 |  | Homo sapiens | leukemia | GPL26945 | 48 |
| GSE131790 |  | Homo sapiens | ovarian cancer | GPL8786 | 25 |
| GSE143564 | 32175269 | Homo sapiens | breast cancer | GPL21572 | 6 |
| GSE124566 | 31938073 | Homo sapiens | head and neck cancer | GPL18402 | 20 |
| GSE144463 |  | Homo sapiens | breast cancer | GPL15468 | 50 |
| GSE134266 | 33391510 | Homo sapiens | prostate cancer | GPL25143 | 29 |
| GSE140719 | 32232006 | Homo sapiens | pancreatic cancer | GPL19117 | 11 |
| GSE135918 | 32194819 | Homo sapiens | lung cancer | GPL18058 | 10 |
| GSE148846 | 32509578 | Homo sapiens | breast cancer | GPL21576 | 24 |
| GSE100682 |  | Homo sapiens | hemangioma | GPL21572 | 6 |
| GSE147889 | 33303811 | Homo sapiens | hepatocellular carcinoma | GPL21263 | 194 |
| GSE100775 |  | Homo sapiens | brain cancer | GPL23653 | 6 |
| GSE116994 |  | Homo sapiens | larynx cancer | GPL19117 | 10 |
| GSE123377 | 32393661 | Homo sapiens | pancreatic cancer | GPL19117 | 17 |
| GSE155467 |  | Homo sapiens | breast cancer | GPL21572 | 6 |
| GSE140001 | 32710569 | Homo sapiens | biliary tract cancer | GPL24158 | 36 |
| GSE143385 | 33148256 | Homo sapiens | adrenocortical cancer | GPL27982 | 39 |
| GSE103996 | 31906302 | Homo sapiens | thyroid cancer | GPL20194 | 34 |
| GSE152702 | 32726984 | Homo sapiens | lung cancer | GPL24158 | 45 |
| GSE158284 |  | Homo sapiens | brain cancer | GPL13264 | 41 |
| GSE159959 |  | Homo sapiens | uterus cancer | GPL29277 | 12 |
| GSE159979 | 33324567 | Homo sapiens | breast cancer | GPL21572 | 9 |
| GSE153089 | 33282720 | Homo sapiens | hepatocellular carcinoma | GPL21572 | 44 |
| GSE163031 |  | Homo sapiens | pancreatic cancer | GPL19117 | 15 |
| GSE139031 | 31808923 | Homo sapiens | brain cancer, lymphoma | GPL21263 | 414 |
| GSE122497 | 31125107 | Homo sapiens | esophageal cancer | GPL21263 | 5531 |
| GSE113740 | 32025611 | Homo sapiens | breast cancer, colorectal cancer, esophageal cancer, gastric cancer, hepatocellular carcinoma, lung cancer, sarcoma, ovarian cancer, pancreatic cancer, brain cancer, prostate cancer, bladder cancer, biliary tract cancer | GPL21263 | 1817 |
| GSE112264 | 30808771 | Homo sapiens | colorectal cancer, esophageal cancer, gastric cancer, hepatocellular carcinoma, lung cancer, sarcoma, pancreatic cancer, brain cancer, prostate cancer, bladder cancer, biliary tract cancer | GPL21263 | 1591 |
| E_MTAB_4667 |  | Homo sapiens | ovarian cancer |  | 125 |
| E_MTAB_4490 | 27625077 | Homo sapiens | gastrointestinal stromal tumor |  | 13 |
| E_MTAB_4170 | 27137948 | Homo sapiens | hepatocellular carcinoma | GPL18941 | 48 |
| E_MTAB_3397 | 26020509 | Homo sapiens | prostate cancer | GPL7731 | 193 |
| E_MTAB_3273 | 26250552 | Homo sapiens | sarcoma |  | 10 |
| E_MTAB_2479 | 25143946 | Homo sapiens | colorectal cancer |  | 104 |
| E_MTAB_1454 |  | Homo sapiens | leukemia |  | 233 |
| E_MTAB_1067 | 24803669 | Homo sapiens | ovarian cancer |  | 243 |
| E_MTAB_862 |  | Homo sapiens | small intestinal neuroendocrine tumor |  | 9 |
| E_MTAB_736 | 22049245 | Homo sapiens | thyroid cancer |  | 22 |
| E_MTAB_408 | 22266859 | Homo sapiens | prostate cancer |  | 54 |
| E_MTAB_386 | 22348002 | Homo sapiens | ovarian cancer |  | 115 |
| GSE39833 | 24705249 | Homo sapiens | colorectal cancer | GPL14767 | 99 |
| GSE85677 |  | Homo sapiens | hepatocellular carcinoma | GPL21263 | 103 |
| GSE76449 | 27742688 | Homo sapiens | ovarian cancer | GPL19117 | 26 |
| GSE125030 |  | Homo sapiens | melanoma | GPL19117 | 10 |
| GSE159177 | 32191585 | Homo sapiens | prostate cancer | GPL21572 | 344 |
| GSE138092 |  | Homo sapiens | brain cancer | GPL19117 | 65 |
| GSE125191 | 32328453 | Homo sapiens | brain cancer | GPL25134 | 20 |
| GSE97135 | 28961307 | Homo sapiens | leukemia | GPL21572 | 39 |
| GSE88958 |  | Homo sapiens | prostate cancer | GPL14767 | 40 |
| GSE125961 | 32293320 | Homo sapiens | colorectal cancer | miRNA-seq | 12 |
| GSE125904 | 30895943 | Homo sapiens | colorectal cancer | miRNA-seq | 8 |
| DRP001085 | 24520312 | Homo sapiens | bladder cancer | miRNA-seq | 10 |
| GSE69810 | 29242506 | Homo sapiens | lymphoma | miRNA-seq | 13 |
| GSE72080 | 26355751 | Homo sapiens | breast cancer | miRNA-seq | 32 |
| GSE57381 | 25567797 | Homo sapiens | hepatocellular carcinoma | miRNA-seq | 12 |
| SRP040720 |  | Homo sapiens | lung cancer | miRNA-seq | 19 |
| GSE67004 | 26132860 | Homo sapiens | colorectal cancer | miRNA-seq | 18 |
| GSE152167 | 32445454 | Homo sapiens | lung cancer | miRNA-seq | 6 |
| SRP091416 |  | Homo sapiens | lymphoma | miRNA-seq | 6 |
| GSE84306 | 29137278 | Homo sapiens | head and neck cancer | miRNA-seq | 15 |
| GSE63046 | 24875649 | Homo sapiens | hepatocellular carcinoma | miRNA-seq | 48 |
| GSE39162 | 21199797 | Homo sapiens | breast cancer | miRNA-seq | 10 |
| GSE130512 | 32508877 | Homo sapiens | thyroid cancer | miRNA-seq | 24 |
| GSE117452 | 30321299 | Homo sapiens | breast cancer | miRNA-seq | 68 |
| SRP132656 |  | Homo sapiens | colorectal cancer | miRNA-seq | 100 |
| GSE68085 | 27177224 | Homo sapiens | breast cancer | miRNA-seq | 114 |
| GSE49279 | 24747642 | Homo sapiens | adrenocortical cancer | miRNA-seq | 48 |
| GSE31616 |  | Homo sapiens | bladder cancer, testicular cancer | miRNA-seq | 34 |
| GSE137308 |  | Homo sapiens | larynx cancer | miRNA-seq | 6 |
| SRP194929 |  | Homo sapiens | colon cancer | miRNA-seq | 25 |
| GSE66186 | 25971364 | Homo sapiens | leukemia | miRNA-seq | 70 |
| GSE63511 | 26282166 | Homo sapiens | thyroid cancer | miRNA-seq | 23 |
| ERP120483 |  | Homo sapiens | breast cancer | miRNA-seq | 54 |
| GSE150956 | 32630542 | Homo sapiens | brain cancer | miRNA-seq | 273 |
| GSE158659 | 33173155 | Homo sapiens | cervical cancer, head and neck cancer | miRNA-seq | 42 |
| GSE10891 | 18538733 | Mus musculus | lymphoma | GPL6606 | 12 |
| GSE12394 | 18713946 | Mus musculus | lymphoma | GPL7152 | 54 |
| GSE23977 | 21846369 | Mus musculus | breast cancer | GPL10880 | 11 |
| GSE23978 | 21846369 | Mus musculus | breast cancer | GPL10880 | 46 |
| GSE29314 | 22052531 | Mus musculus | prostate cancer | GPL8786 | 6 |
| GSE34738 | 23479506 | Rattus norvegicus | pancreatic cancer | GPL7723 | 8 |
| GSE35402 | 22733778 | Mus musculus | liver cancer | GPL15169 | 6 |
| GSE41723 | 23509284 | Mus musculus | breast cancer | GPL16199 | 15 |
| GSE44983 | 23615396 | Mus musculus | colorectal cancer | GPL16759 | 6 |
| GSE44570 |  | Mus musculus | hepatocellular carcinoma | GPL16713 | 16 |
| GSE50748 | 24265725 | Mus musculus | lung cancer | GPL8786 | 18 |
| GSE50752 | 24265725 | Mus musculus | lung cancer | GPL13493 | 24 |
| GSE47211 | 24349037 | Mus musculus | melanoma | GPL16275 | 13 |
| GSE54177 | 24735923 | Mus musculus | colorectal cancer | GPL17873 | 18 |
| GSE56759 | 24735967 | Mus musculus | lymphoma | GPL17568 | 6 |
| GSE43483 | 25378704 | Mus musculus | breast cancer | GPL7732 | 12 |
| GSE75862 | 27729023 | Mus musculus | lung cancer | GPL7732 | 6 |
| GSE63633 | 26059540 | Mus musculus | breast cancer | GPL19460 | 15 |
| GSE84109 | 28599420 | Mus musculus | lung cancer | GPL7732 | 6 |
| GSE68630 | 26749280 | Mus musculus | breast cancer | GPL16384 | 9 |
| GSE98391 | 29196616 | Mus musculus | ovarian cancer | GPL21572 | 6 |
| GSE93936 |  | Mus musculus | colon cancer | GPL19117 | 10 |
| GSE132052 |  | Mus musculus | brain cancer | GPL26722 | 7 |
| GSE102883 |  | Mus musculus | melanoma | GPL14613 | 8 |
| GSE126549 | 31503420 | Mus musculus | ovarian cancer | GPL22601 | 17 |
| GSE139252 | 32306082 | Mus musculus | hepatocellular carcinoma | GPL16384 | 15 |
| GSE145083 | 33407852 | Mus musculus | lymphoma | GPL19117 | 8 |
| GSE162793 | 33652981 | Mus musculus | breast cancer | GPL29478 | 7 |
| GSE34119 | 23273170 | Rattus norvegicus | breast cancer | GPL10344 | 23 |
| GSE156430 | 32842712 | Mus musculus | colorectal cancer | GPL16199 | 6 |
| GSE114110 | 30361064 | Homo sapiens | esophageal cancer | GPL24967 | 40 |
| GSE86100 |  | Homo sapiens | cervical cancer | GPL19730 | 12 |
| GSE135189 | 30570705 | Homo sapiens | brain cancer | GPL20906 | 28 |
| GSE131166 | 32244823 | Homo sapiens | head and neck cancer | GPL16770 | 6 |
| GSE134108 | 31603918 | Homo sapiens | breast cancer | GPL18941 | 8 |
| GSE140584 |  | Homo sapiens | bladder cancer | GPL27765 | 6 |
