## Supplementary material for "dbDEMC 3.0: Functional Exploration of Differentially Expressed miRNAs in Cancers of Human and Model Organisms": Table S3

**Table S3 Cancer types and associated subtypes and cell line names covered in dbDEMC 3.0.**

| Cancer Type | Cancer Subtypes | Cell Line |
| --- | --- | --- |
| adrenocortical cancer | adrenocortical carcinoma |  |
| biliary tract cancer | cholangiocarcinoma, cholangiocellular carcinoma, distal cholangiocarcinoma |  |
| bladder cancer | bladder urothelial carcinoma |  |
| brain cancer | medulloblastoma, glioblastoma, meningioma, Schwannoma tumors, glioma, anaplastic astrocytoma, glioblastoma multiforme, brainstem gliomas, astrocytoma, pilocytic astrocytoma |  |
| breast cancer | ER positive, ER negative, Luminal A, Luminal B, HER2+, Basal-like, Normal-like, PR positive, PR negative, invasive ductal carcinoma, triple negative, invasive lobular carcinoma, squamous cell breast carcinoma, acantholytic variant | MCF-7, LCC9, LCC2, 67NR, 168FARN, 4TO7, 4T1 |
| cervical cancer | cervical adenocarcinoma, cervical squamous cell carcinoma |  |
| chordoma |  |  |
| colon cancer | colonic adenomas, tubulovillous adenoma, serrated adenoma, colon adenocarcinoma | SW620, LoVo, FHC |
| colorectal cancer | Duke A, Duke B, Duke C, colorectal adenocarcinoma | HCT116, DKO-1, Dks-8, DLD-1 |
| endometrial cancer | uterine corpus endometrial carcinoma |  |
| esophageal cancer | Barrett's carcinogenesis, esophageal squamous cell carcinomas, esophageal adenocarcinoma | OE19 |
| gallbladder carcinoma |  |  |
| gastric cancer | H. Pylori positive gastric cancer, H. Pylori negative gastric cancer |  |
| gastrointestinal stromal tumor |  |  |
| head and neck cancer | HPV positive, HPV negative, head and neck squamous cell carcinoma | Cal 27, H413, HGEPp, Detroit 562, FaDu |
| hemangioma | infantile hemangioma |  |
| hepatocellular carcinoma | hepatocellular carcinoma, hepatoblastoma | HepG2 |
| kidney cancer | Wilms tumor, clear cell renal cell carcinoma, papillary renal cell carcinoma, chromophobe renal cell carcinoma, Oncocytoma, kidney renal clear cell carcinoma, kidney chromophobe cancer |  |
| larynx cancer | aryngeal squamous cell carcinoma |  |
| leukemia | T cell leukemia, acute lymphocytic leukemia, acute myelogenous leukemia, B acute lymphoblastic leukemia, T acute lymphoblastic leukemia, acute myelocytic leukemia, chronic lymphocytic leukemia, chronic myeloid leukemia, plasma cell leukemia | Jurkat |
| liver cancer | intrahepatic cholangiocarcinoma, combined hepatocellular cholangiocarcinoma, cholangiocarcinoma |  |
| lung cancer | lung squamous cell carcinoma, lung adenocarcinoma, small cell lung cancer, large cell neuroendocrine cancer, large cell carcinoma, non-small cell lung cancer | H446, LLC1 |
| lymphoma | B cell lymphoma, Burkitt's lymphoma, Splenic lymphoma, Gastric lymphoma, Lymphoblastoid, follicular cleaved lymphoma, diffuse large B cell lymphoma, multiple myeloma, B cell lymphoma of MALT type, B-cell non-Hodgkin lymphoma, Hodgkin lymphoma, myeloma, T-cell lymphoblastic lymphoma, primary central nervous system lymphomas, diffuse large B-cell lymphoma, MYC-dependent lymphoma, CD4+ T cell lymphomas, chronic myelomonocytic leukemia | OCI-Ly4, OCI-Ly7, OCI-Ly8, Namalwa, Raji, Karpas-1718, Manca, HG-1125, H929 |
| melanoma | skin cutaneous melanoma, melanoma brain metastasis | A375, A375R, M117,M113, B16F0 |
| mesothelioma | malignant pleural mesothelioma, diffuse malignant peritoneal mesothelioma |  |
| nasopharyngeal cancer |  |  |
| neuroendocrine neoplasia |  |  |
| oral squamous cell carcinoma |  |  |
| oropharyngeal squamous cell carcinoma |  |  |
| ovarian cancer | clear cell ovarian carcinoma, high-grade serous ovarian carcinoma, familial ovarian cancer, high grade serous papillary ovarian cancer, serous ovarian cancers | HIO180, SKOV3_ip1, SKOV3_TR, HEYA8, A2780 |
| pancreatic cancer | pancreatic ductal adenocarcinoma, Solid-pseudopapillary neoplasm of pancreas, pancreatic adenocarcinoma, solid pseudopapillary neoplasm of pancreas | PaCa-2, S2-013, PANC-1, ASML |
| prostate cancer | prostate adenocarcinoma, castration-resistant prostate cancer |  |
| retinoblastoma |  |  |
| sarcoma | Synovial sarcoma, liposarcoma, osteosarcoma, Kaposi's sarcoma, rhabdomyosarcomas, synovial sarcoma |  |
| skin cancer |  |  |
| small intestinal neuroendocrine tumor |  |  |
| testicular cancer | testicular germ cell tumor |  |
| thyroid cancer | primary papillary thyroid carcinomas, medullary thyroid cancer, papillary thyroid carcinoma, follicular thyroid carcinoma |  |
| tonsil cancer |  |  |
| uterus cancer |  |  |
